# Single-Cell Proteomic and Transcriptomic Characterization of Drug-Resistant Prostate Cancer Cells Reveals Molecular Signatures Associated with Morphological Changes

**DOI:** 10.1101/2024.10.23.619905

**Authors:** Jongmin Woo, Michael Loycano, Md Amanullah, Jiang Qian, Sarah Amend, Kenneth Pienta, Hui Zhang

## Abstract

This study delves into the proteomic intricacies of drug-resistant cells (DRCs) within prostate cancer, which are known for their pivotal roles in therapeutic resistance, relapse, and metastasis. Utilizing single-cell proteomics (SCP) with an optimized high-throughput Data Independent Acquisition (DIA) approach with the throughput of 60 sample per day, we characterized the proteomic landscape of DRCs in comparison to parental PC3 cells. This optimized DIA method allowed for robust and reproducible protein quantification at the single-cell level, enabling the identification and quantification of over 1,300 proteins per cell on average. Distinct proteomic sub-clusters within the DRC population were identified, closely linked to variations in cell size. The study uncovered novel protein signatures, including the regulation of proteins critical for cell adhesion and metabolic processes, as well as the upregulation of surface proteins and transcription factors pivotal for cancer progression. Furthermore, by integrating SCP and single-cell RNA-seq (scRNA-seq) data, we identified six upregulated and ten downregulated genes consistently altered in drug-treated cells across both SCP and scRNA-seq platforms. These findings underscore the heterogeneity of DRCs and their unique molecular signatures, providing valuable insights into their biological behavior and potential therapeutic targets.

## Introduction

Cancer is a heterogeneous disease characterized by diverse cell populations with distinct genetic and phenotypic profiles (1, 2). Among these, drug-resistant cells (DRCs) have garnered significant attention due to their association with therapeutic resistance, disease relapse, and metastasis (3, 4). DRCs are characterized by their large size and complex karyotypes, which contribute to their adaptability and survival under therapeutic stress (5, 6). DRCs have been identified in various cancer types, including breast, colorectal, and ovarian cancers, where they correlate with poor prognosis and survival (7-12).

These cells often arise due to oncogenic and therapeutic stress, leading to genomic instability and the formation of cells with multiple nuclei and enlarged size (13, 14). Recent advancements in single-cell technologies have enabled detailed molecular characterizations of these rare cell populations (11). While single-cell RNA-seq (scRNA-seq) has provided valuable insights into the transcriptional landscapes of DRCs, proteomics offers a complementary approach by directly measuring the functional protein molecules that drive cellular behavior (15, 16). Therefore, this study focuses on proteomic characterization of DRCs of prostate cancer cells in single-cell level, aiming to uncover the proteomic alterations that underpin their unique biological properties.

Single-cell proteomics, particularly when combined with high-throughput methodologies such as Data Independent Acquisition Mass Spectrometry (DIA-MS) (17) and Liquid Chromatography with Field Asymmetric Ion Mobility Spectrometry Mass Spectrometry (LC-FAIMS-MS) (18), allows for an in-depth analysis of the proteomic landscape of these cells. By employing state-of-the-art single-cell proteomics, we not only capture the extensive proteomic repertoire of these cells but also benchmark the reproducibility and quantification capabilities for single-cell proteomic characterization of these cells.

This benchmarking is critical, as it ensures the reliability of data especially when comparing the complex proteomes of distinct cell populations like DRCs and their parental cell lines. In addition, isolating DRCs poses significant challenges due to their size (19, 20). In this study, we employed the CellenOne instrument for cell isolation, addressing size-related issues by diluting and percolating cells through a strainer to ensure accurate sampling. This study marks the first detailed molecular characterization of DRCs at the single-cell level, providing novel insights into their biology and potential therapeutic vulnerabilities. By integrating these findings with scRNA-seq, we aim to contribute to the growing body of knowledge on DRCs and highlight the potential of single-cell proteomics in advancing clinical cancer research. Our work underscores the importance of studying these enigmatic cells to develop more effective therapeutic strategies against prostate cancer

## Methods

### Cell culture and drug-resistance cell induction

Cell culture experiments were conducted using the PC3-luciferase-GFP prostate cancer cell line (ATCC and edited by the UMich Cell Line Core and Pienta’s group at JHU). The cells were cultured in RPMI 1640 medium with L-glutamine and phenol red (Gibco), supplemented with 10% Premium Grade Fetal Bovine Serum (Avantor Seradigm) and 1% Penicillin-Streptomycin (5000 U/mL; Gibco), and maintained at 37°C in a 5% CO2 incubator. Authentication and mycoplasma contamination testing of the PC3 cells were performed biannually by Genetica. Cells were seeded at a density of 6,000,000 cells per 500cm2 dish (Corning). Twenty-four hours post-seeding, the cells were treated with 6 μM (GI50) cisplatin, dissolved in PBS with 140 mM NaCl, for 72 hours (Millipore Sigma). After the treatment, the medium was replaced with fresh medium, and the cells were cultured for an additional 10 days, at which point they were considered definitive drug-resistant cells (DRCs). The medium was refreshed every 4 days.

TrypLE Express Enzyme without phenol red was routinely used for cell dissociation and plates were scraped using a cell scraper. All centrifugation was done at 200 xg for 10 minutes. Cells were resuspended in PBS for transport. Images were captured using the EVOS M7000 and EVOS FL Auto microscopes (ThermoFisher, Figure 1). Both PC3 parental and DRCs were immediately transported on ice for cell isolation.

**Figure 1.**
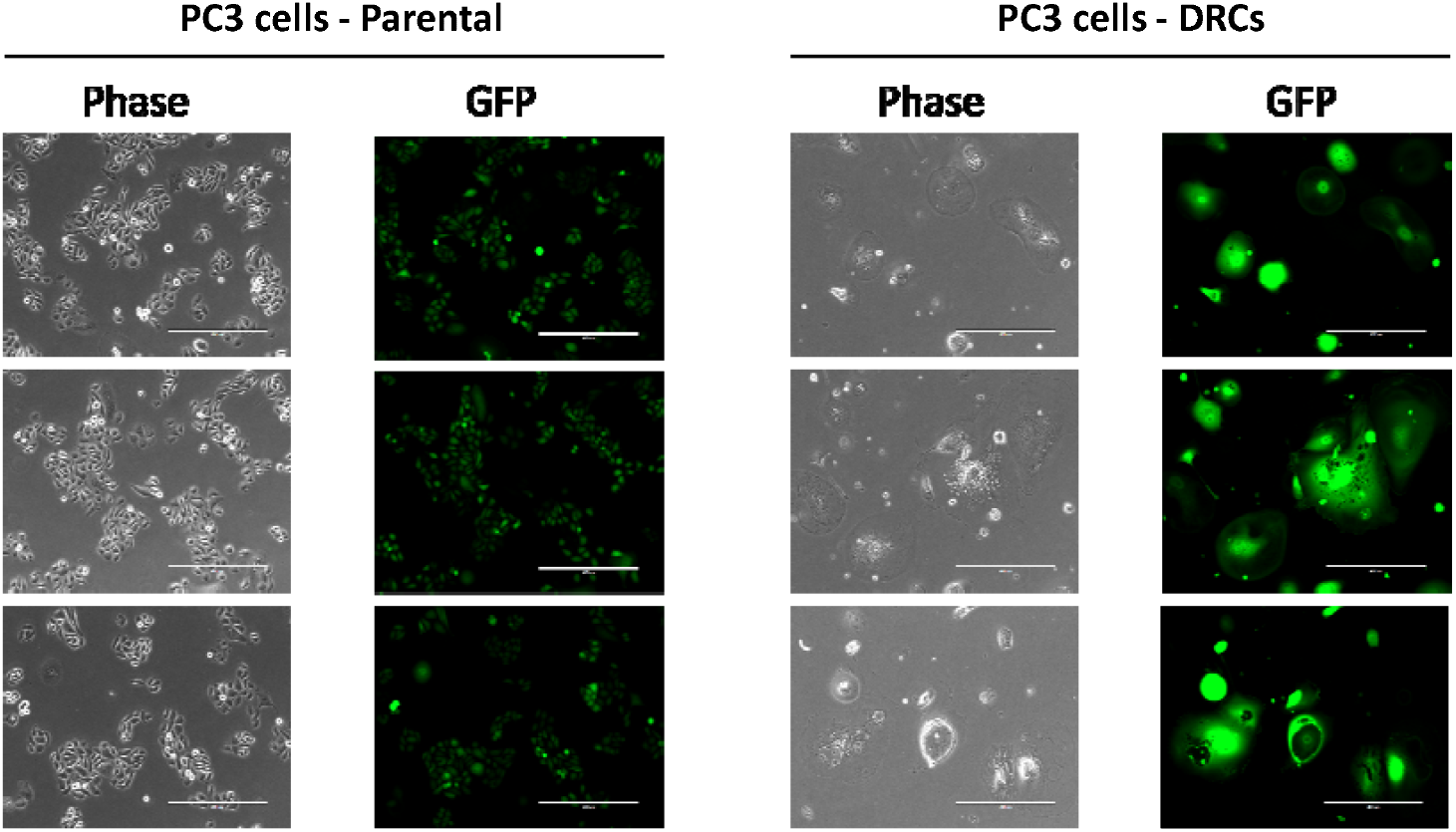
Morphological and fluorescence analysis of DRCs and Parental PC3 cells. Left panels in each group present bright-field images demonstrating the morphological characteristics of parental PC3 cells and DRCs. Right panels show corresponding fluorescence images.

### Single-cell isolation for proteomics and RNA-seq studies

Single cells were isolated using the CellenOne instrument, which enables precise manipulation of individual cells via real-time imaging that assesses cell size. PC3 parental cells were first diluted in PBS to a concentration of 200 cells/µL and isolated based on a cell size range of 22 to 25 µm in diameter. For the DRCs, the dilution was adjusted to 50 cells/µL to prevent clogging of the nozzle caused by the larger cell sizes. DRCs were then isolated based on a cell size range of 45 to 102 µm in diameter. Each single cell was dispensed into a well of a 384-well plate containing 1 µL of pre-added lysis buffer, prepared for the subsequent digestion procedure. During the isolation process, images of all isolated cells were captured, including measurements of cell diameter and bright-field intensity, for further analysis.

### Single-cell sample preparation for proteomic analysis

Isolated cells underwent lysis, protein extraction, and digestion, as the on-plate sample preparation (21). In brief, 1 μL of lysis buffer—comprising 0.1% DDM, 0.02% LMNG, and 5 mM TCEP in a 100 mM TEAB buffer—was added to each well of a 384-well plate prior to cell isolation. After cell deposition, the plate was centrifuged briefly at 1000 x g for 5 minutes. Subsequently, 5 ng of a Trypsin/LysC mixture was added to each well, followed by overnight incubation at 37°C. The resulting peptide samples were acidified and stored at -80°C until ready for LC-MS analysis. For loading onto the Evotips, which had been washed and activated according to the manufacturer’s instructions, peptides were first thoroughly washed in 10 μL of 0.1% formic acid before being transferred to the Evotips.

### Serial diluted peptide sample preparation

NCI-7 cells were used as reference materials (David Clark, ref). Peptides from NCI-7 cells were prepared following the CPTAC protocol (22). Briefly, 1×10^6 cells were lysed via centrifugation at 10,000 x g for 10 minutes. The extracted proteins were denatured using 8M urea, reduced with 10 mM DTT, and alkylated with 20 mM IAA. The proteins were then digested with trypsin and the resulting peptides were serially diluted across nine dilution points, ranging from 400 ng to 125 pg.

### Proteomic analysis using LC-MS instruments

Proteomic analyses were performed by DIA-MS using two distinct instruments paired with the Evosep LC: the TimsTOF HT (Bruker) and the Orbitrap Ascend (Thermo). Peptide samples were separated in the Evosep using short gradients, allowing for high-throughput analysis of up to 100 samples per day (SPD) utilizing an analytical column (EV1064, 8 cm x 100 μm x 3 μm). The trapped ion mobility spectrometry (Tims) Parameters for the TimsTOF included a mobility range of 0.70 – 1.36 1/k0, mass range of ?, and an accumulation time of 166 ms with high-sensitivity detection enabled. The Orbitrap Ascend utilizes

Field Asymmetric Ion Mobility Spectrometry (FAIMS) FAIMS and the settings were adjusted to a compensational voltage of -55 V, a mass range of 380-980 m/z, an isolation window of 35 Th, a resolution of 30k, an Automatic Gain Control (AGC) of 2.5×10E5, and an maximum injection time (IT) of 166 ms.

In the single-cell analysis using the Orbitrap Ascend coupled with Evosep LC and FAIMS (−55 CV), single-cell samples were separated utilizing a throughput of 60 SPD with the same analytical column. Parameters remained consistent with those listed above, but the mass range was adjusted to 380-780 m/z and the isolation window was narrowed to 25 Th to optimize protein identification and quantification.

### Data Analysis for scProteomics

Proteomic raw data from TimsTOF HT and Ascend were processed using Spectronaut (Ver 18.0) with the directDIA approach to benchmark the two datasets. Single-cell data obtained from FAIMS-Ascend were analyzed using DIA-NN (Ver 1.8.0) employing a library-based database search. This library was created by injecting 1 ng of peptide samples from either PC3-parental or DRC, prepared via bulk sample prep, containing 7095 precursors and 2034 proteins. In DIA-NN, both MS1 and MS2 accuracies were set at 10 ppm, with a scan window of 3. Features such as using isotopologues, match-between-runs (MBR), and unrelated runs were enabled. Cross-normalization was disabled. After log2 transformation of the quantitative values, protein groups were recognized as identified if they had quantitative values, and as quantified if at least 70% of the samples in each group presented valid quantitative measurements, resulting in improving data imputation accuracy and increasing statistical power. The missing values were imputed using KNN with a neighbor number of 15. Statistical methods such as Principal Component Analysis (PCA) and Uniform Manifold Approximation and Projection (UMAP), combined with k-means clustering, were used to identify distinct proteomic profiles and sub-clusters within the DRC population. A two-sample t-test was conducted to determine significant protein regulation, with a significance threshold set at p-value < 0.05. Additionally, correlation and bioinformatics analyses were performed to investigate the relationship between cell volume and proteome abundance.

### Transcriptomics analysis

RNA from single cells in lysis buffer was immediately processed for reverse transcription. RNA quality was assessed using an Agilent Bioanalyzer (Agilent Technologies, Santa Clara, CA, USA). mRNA reverse transcription was performed using the SMART-Seq v4 Oligonucleotide technology, which employs oligo(dT) priming to synthesize full-length cDNA. The reaction was carried out at 42°C for 90 minutes, followed by enzyme inactivation at 70°C for 10 minutes. The first-strand cDNA synthesis was optimized with the inclusion of locked nucleic acids (LNA) to enhance template-switching efficiency. Amplification of the first-strand cDNA was performed using Long-Distance PCR (LD-PCR) with SeqAmp DNA polymerase. The number of PCR cycles was adjusted according to the RNA input, with 17 cycles used for 10 pg of RNA. The amplified cDNA was then purified using AMPure XP beads (Beckman Coulter, USA). cDNA was quantified using the Qubit dsDNA HS Assay (Thermo Fisher Scientific) and assessed for quality using an Agilent Bioanalyzer to confirm the presence of high-quality, full-length cDNA. For sequencing, cDNA libraries were prepared using Takara Bio’s SMART-Seq Library Prep Kit, which utilizes stem-loop adapters for enzymatic fragmentation and indexing. The libraries were further amplified through an additional 12–16 PCR cycles, depending on the input cDNA concentration. After amplification, samples were pooled, purified with AMPure XP beads, and validated for library size distribution using an Agilent Bioanalyzer. Library quantification was conducted via qPCR before sequencing on an Illumina platform, utilizing paired-end 150 bp reads.

### Sequence alignment and gene expression quantification

FastQC (v0.12.1) tool was used to assess the quality of raw sequencing data. The quality of sequencing data demonstrated high-quality metrics, with majority of reads exhibiting Phred quality scores above 30 across all bases. Key quality indicators, such as base sequence quality, GC content, and adapter contamination, were within acceptable ranges, indicating that the sequencing data was of good quality. Paired-end sequencing reads were then mapped to the reference genome using the STAR (2.7.11b) Aligner following the standard pipeline (23). To obtain accurate and high-speed quantification of read counts, featureCounts function from the Subread package (v2.0.6) was used based on the GRCm39.111 GTF annotation data (24). The human reference genome (GRCh38) and gene annotation (GRCh38.111) GTF data were downloaded from Ensemble database. To reduce noise from lowly expressed genes, genes with less than 1 CPM value across the samples were then filtered, leaving 25,651 genes for subsequent analysis. Following this filtering step, only genes with reliable expression patterns were retained for further analysis. Similar to our analysis of the SCP data, quantitative values were log2-transformed.

Protein groups were considered identified if quantitative values were present, and considered quantified if at least 70% of the samples in each group contained valid quantitative measurements. For the comparison between SCP and scRNA-seq data, both datasets were normalized using width adjustment, aligning the median to zero across all quantitative protein groups. Statistical analysis was performed using a t-test with a p-value threshold of <0.05.

## Results

### Evaluation of analytical performances of DIA-MS in proteomic analysis of single-cell-level peptides with two MS instruments

In our study, we initially assessed identified and quantified proteins from serially diluted peptide samples using two different mass spectrometry instruments: the Orbitrap Ascend and the TimsTOF HT. The Ascend utilizes FAIMS, which selectively filters ions based on their differential mobility in an oscillating electric field. This selective ion passage significantly reduces chemical noise and boosts the signal-to-noise ratio, which is critical for detecting low-abundance peptides in low-input samples, such as those typical in single-cell proteomics (18). Conversely, the timsTOF HT employs PASEF-DIA, incorporating Tims followed by time-of-flight (TOF) mass spectrometry. TimsTOF is effective to achieve accumulation of ions (Tims) and increased instrument speed and throughput (TOF) (25). Furthermore, the Ascend features Automatic Gain Control (AGC), optimizing the ion count entering the mass spectrometer to ensure stable and consistent signals. AGC dynamically adjusts the ion influx, preventing the detector from being underutilized or overwhelmed, thus enhancing the accuracy and reliability of measurements in samples with extremely low ion counts. In contrast, the timsTOF HT uses ion mobility for ion capture.

However, both instruments demonstrated comparable results, as illustrated in Figures 2A, 2B, and 2C, increased number of proteins were identified and quantified with high-input peptide amount (>=50 ng) for TimsTOF and elevated number of proteins were identified and quantified with low sample input for Oritrap Ascend. Based on our evaluation, we decided to use FAIMS-DIA on the Ascend for further single-cell analysis due to its superior performance in handling low-input samples and its enhanced measurement reliability.

**Figure 2.**
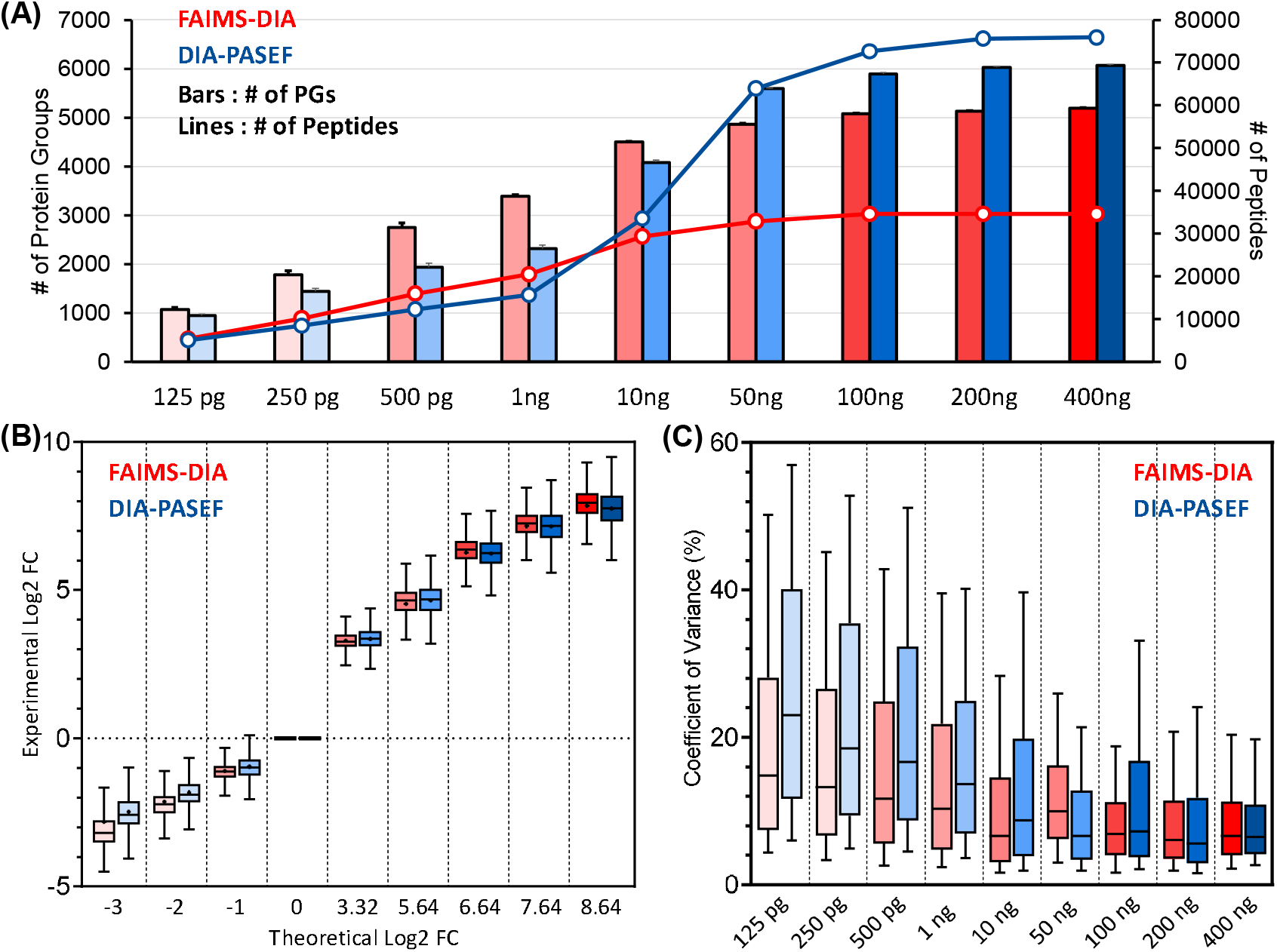
Analytical performances of DIA-MS in proteomic analysis of single-cell-level peptides with two MS instruments. (A) Identified peptides and protein groups from serially diluted NCI-7 peptide samples. In this figure, red indicates FAIMS-DIA-MS data from Orbitrap Ascend and blue presented DIA-PASEF data from TimsTOF HT instrument. (B) Linearity of quantification using the two instruments. Each box-plot presents the distribution of Log2 fold-changes average value of protein groups in the sample to the protein group in 1 ng sample. X-axis shows the theoretical Log2 fold-change we can expect. (C) Reproducibility in each sample per dataset from the two different instruments.

### Comparison of proteome abundance and cell volume between parental PC3 cells and DRCs

Given the larger size of DRCs, traditional single-cell isolation methods proved challenging. Initially, we utilized the CellenOne instrument with a recommended cell dilution of 200 cells/μL, but frequently found that it resulted in nozzle clogging during cell sorting. To address this, we implemented two modifications: 1) increasing the dilution to 50 cells/μL to reduce cell congestion at the nozzle and allow more space for cells to undergo morphological changes; 2) employing a strainer to filter out the cells clustered together as well as extremely large cells that caused nozzle clogging during cell sorting using CellenOne. Although it was not essential to eliminate the large cells for DRCs studies, this adjustment enabled us to continue working with significantly large DRC cells without issues.

Further analysis of cell size distribution showed marked differences between DRCs and parental PC3 cells. Parental PC3 cells were found to have diameters ranging from 22 to 25 µm, whereas DRCs varied between 45 and 102 µm, demonstrating the substantial morphological changes associated with polyaneuploidy in DRCs (Figure 3A). Proteomic analysis revealed an average of 1,174 protein groups per parental PC3 cell and 1,367 protein groups per DRC cell, totaling 1,567 unique proteins identified across both cell types (Figure 3B, Supplementary table S1). Quantification of total protein abundance showed significantly higher levels in DRCs compared to parental cells (Figure 3C), correlating with the increased cell volume observed in DRCs. This suggests that DRCs possess a heightened metabolic and biosynthetic capacity to support their larger size and genomic content. A strong positive correlation (Pearson correlation coefficient of 0.69) was found between cell volume and total protein abundance in DRCs, indicating that larger DRCs have greater proteomic capacity (Figure 3D). This relationship underscores the influence of cell size on the functional capabilities of these cells, highlighting the importance of considering cell volume in proteomic studies of polyaneuploid cells.

**Figure 3.**
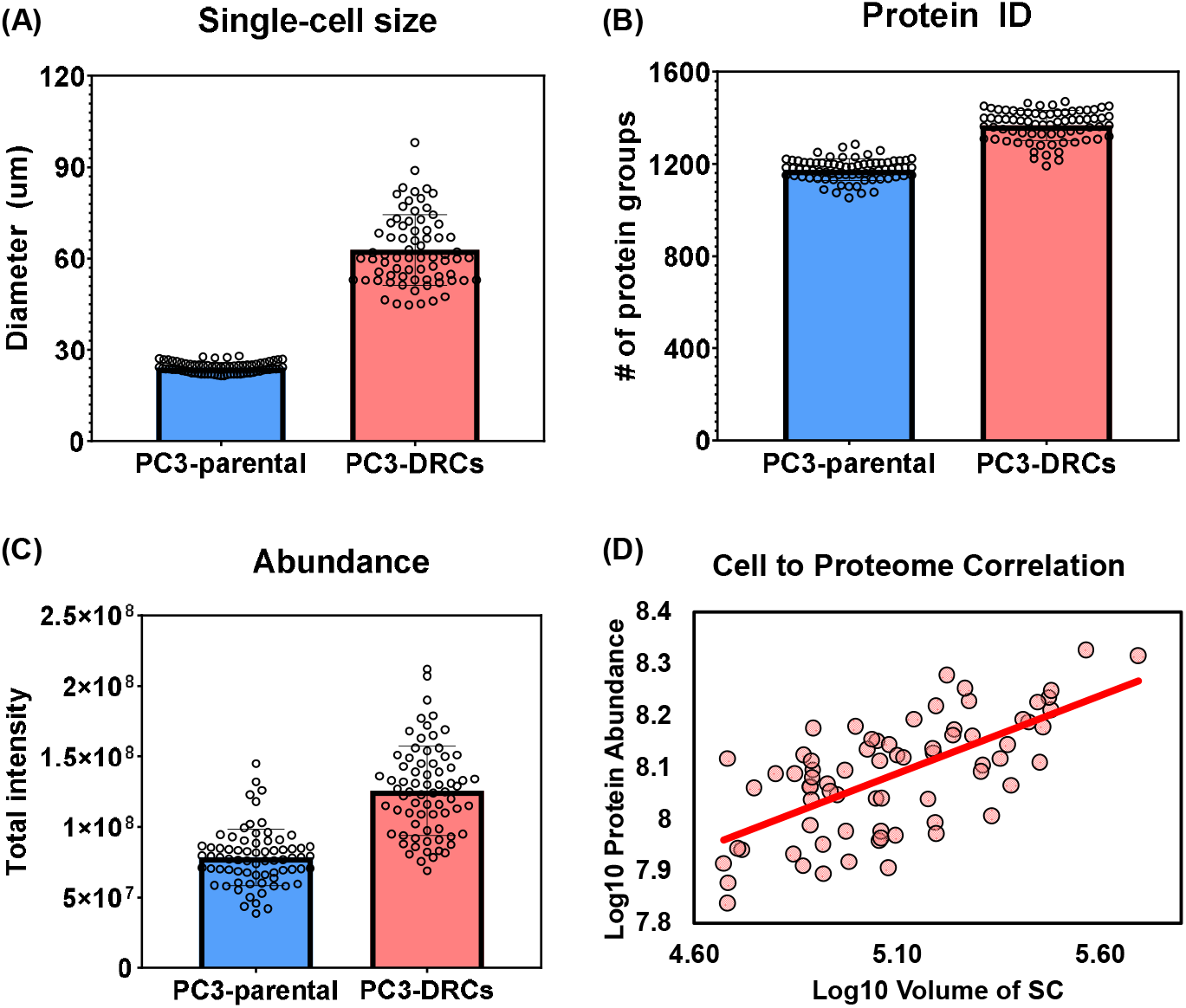
Comparison of proteome abundance and cell volume between parental PC3 cells and PC3-DRCs. (A) Cell size distribution in diameter. (B) Identified protein groups per cell. (C) Total protein abundance per cell. (D) Correlation between cell volume and proteome abundance.

### Multidimensional analysis of single-cell proteome reveals distinct proteomic landscapes and subtype diversity in DRCs of prostate cancer

In our single-cell proteomic analysis, incorporating both parental PC3 cells and DRCs, we employed Principal Component Analysis (PCA) and Uniform Manifold Approximation and Projection (UMAP) to explore underlying proteomic diversities and similarities. The PCA results (Figure 4A) revealed a pronounced segregation between the two cell types, confirming substantial proteomic disparities. This clear demarcation underscored the significant molecular alterations associated with polyaneuploidy in DRCs, which might contribute to their altered physiological states and therapeutic resistance. Further dissecting the proteomic landscape through UMAP, we not only confirmed the distinct clustering between parental PC3 cells and DRCs (Figure 4B) but also delved deeper into the heterogeneity within the DRCs population. The UMAP analysis, supplemented by k-means clustering, discerned three distinct sub-clusters within the DRCs (Figure 4C, Supplementary table S2).

**Figure 4.**
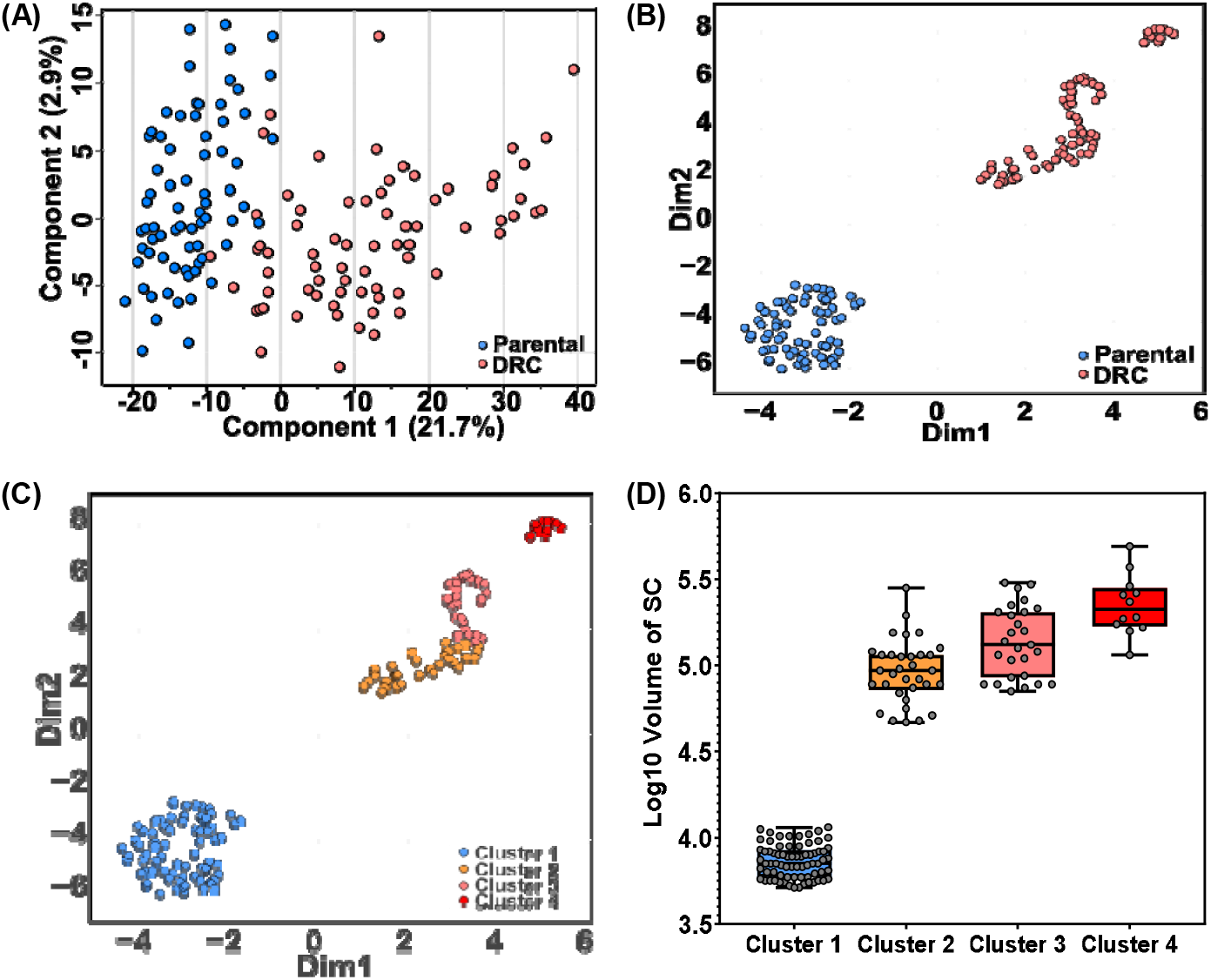
Multidimensional analysis of single-cell proteome reveals distinct proteomic landscapes and subtype diversity in prostate cancer cells, (A) PCA plot of 144 single cells. (B) UMAP - main grouping. (C) UMAP - Three sub-clusters in DRCs using K-means method. (D) Subgroup clusters and cell size association.

The enriched pathways between the clusters reveal key biological themes that distinguish the proteomic profiles of drug-resistant prostate cancer cells. For example, a strong enrichment in mRNA processing (22 observed genes, FDR = 7.36E-08), ribosomal proteins (19 observed genes, FDR = 7.36E-08), and translation factors (11 observed genes, FDR = 0.00012) in significantly regulated proteins in cluster 4 suggests a heightened involvement in protein synthesis and cellular growth, particularly in the clusters associated with more aggressive traits. Additionally, pathways such as VEGFA-VEGFR2 signaling (42 observed genes, FDR = 6.00E-08) and metabolic reprogramming (13 observed genes, FDR = 7.21E-07) highlight shifts in metabolic and signaling processes that support cellular proliferation, survival, and adaptation to therapeutic stress. These results suggest that distinct clusters are not only metabolically reprogrammed but also show enhanced capabilities in protein synthesis and cellular signaling, which likely contribute to their differential responses to treatment and potential for metastasis. This indicates that the proteomic diversity within drug-resistant cells may underpin their heterogeneity in function and survival. Each sub-cluster represents a unique proteomic signature that potentially corresponds to varying degrees of malignancy and resilience against therapeutic interventions. In addition, most compelling is the association of these sub-clusters with cell size variations (Figure 4D), where we observed that larger cell sizes correspond to distinct proteomic profiles. This correlation suggests that cell size, a proxy for cellular complexity and metabolic capacity, might be a critical determinant in the phenotypic and functional heterogeneity observed within DRCs.

### Single-cell proteomic characterization reveals down-regulation of key regulatory proteins in DRCs

In our detailed single-cell proteomic analysis, we identified a significant down-regulation of several proteins in DRCs compared to parental PC3 cells, which could be crucial in understanding the altered physiological states associated with cancer progression and therapy resistance (Figure 5, Supplementary table S3). These proteins, including AHCYL1, APLP2, CTNN1D, EIF2B1, LGALS1, and SGPL1, play vital roles in maintaining cellular functions such as cell adhesion, protein synthesis, and lipid metabolism, which are often perturbed in cancer cells. AHCYL1 (S-adenosylhomocysteine hydrolase-like protein 1) is implicated in the regulation of intracellular homocysteine levels and methylation processes, influencing cell plasticity and potentially tumorigenesis (26); its down-regulation could disrupt these critical metabolic processes, contributing to the aggressive traits of DRCs. APLP2 (Amyloid beta precursor-like protein 2), known for its involvement in synaptic function and cellular adhesion (27), might affect cell-cell interaction and migration dynamics when down-regulated, potentially facilitating cancer cell detachment and metastasis. CTNN1D (Catenin delta-1), a part of the cadherin-catenin complex, plays a significant role in maintaining cell adhesion (28); reduced levels may lead to decreased adhesiveness and increased metastatic potential. EIF2B1 (Eukaryotic translation initiation factor 2B) is essential for the initiation of protein synthesis (29); its reduction could lead to a broad decrease in protein production, affecting various growth and survival pathways in cancer cells. LGALS1 (Galectin-1) functions in cell-matrix interactions and immune response modulation (30); its down-regulation could impact tumor immune evasion and cell communication. SGPL1 (Sphingosine-1-phosphate lyase 1) is critical in sphingolipid metabolism, which is pivotal in regulating apoptosis and cell migration (31); alterations in its levels could modify the balance between cell death and survival, influencing disease progression. The collective down-regulation of these proteins in DRCs underscores a strategic alteration in cellular regulatory mechanisms that could confer survival advantages under therapeutic stress or contribute to an enhanced metastatic capacity. Understanding these protein changes provides valuable insights into the complex molecular dynamics that drive cancer cell adaptation and survival, highlighting potential targets for therapeutic intervention aimed at mitigating the aggressive behaviors of DRCs.

**Figure 5.**
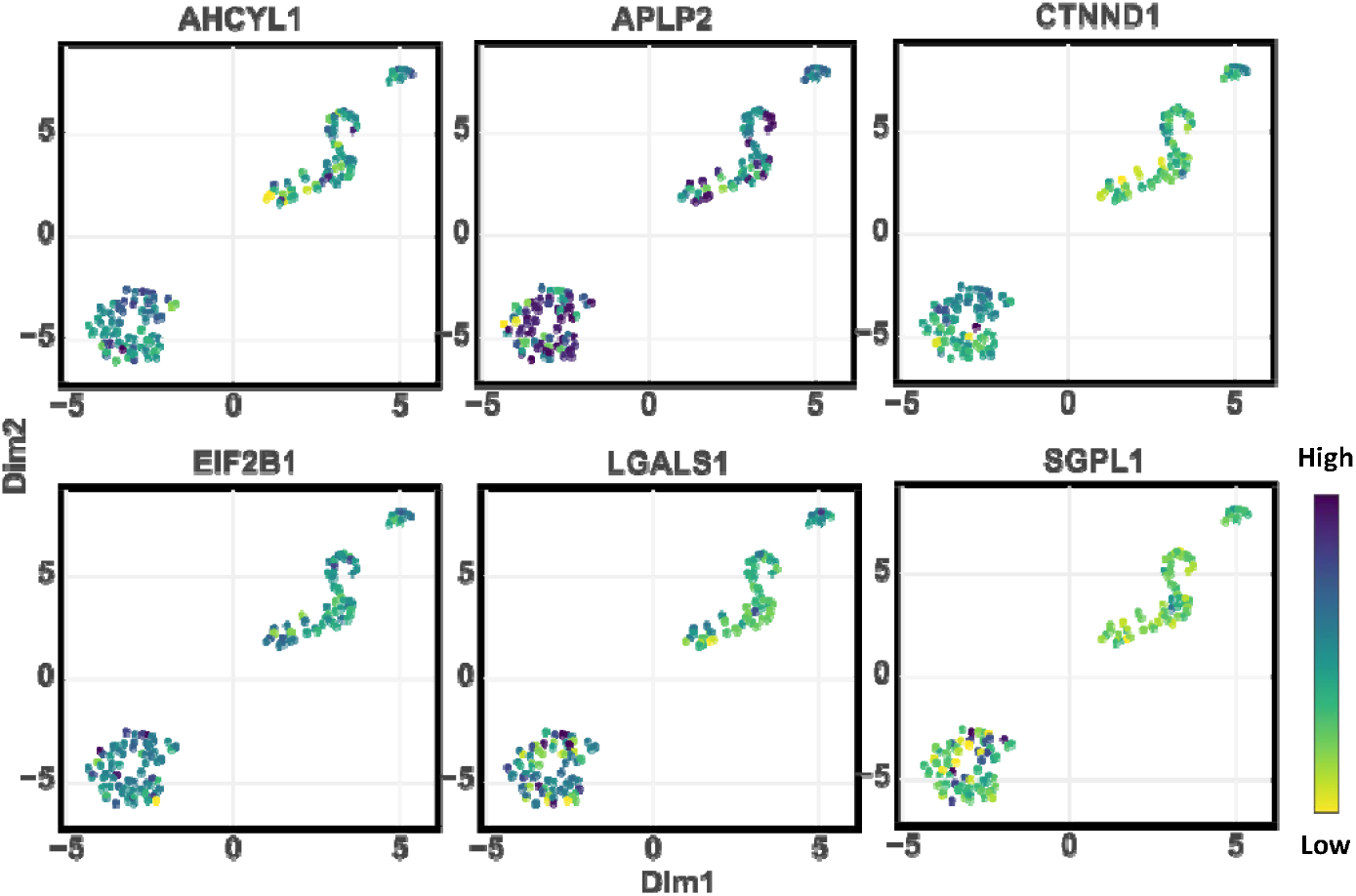
Significant down-regulated proteins in DRCs compared to parental PC3 cells (t-test p-value < 0.05) including AHCYL1, APLP2, CTNN1D, EIF2B1, LGALS1, and SGPL1 were observed.

### Elevated levels of surface proteins and transcription factors underlines DRCs’ aggressive traits

In our analysis, we identified a significant up-regulation of key surface plasma membrane proteins and transcription factors in DRCs, which are likely to play crucial roles in promoting the aggressive behavior and resilience of these cancer cells (Supplementary table S3). Among the surface proteins (Figure 6A), ACTN1 (Alpha-actinin-1), CD47, and FAT1 (FAT atypical cadherin 1) were notably increased. ACTN1, which plays a pivotal role in cell motility and structural integrity (33), might enhance the migratory and invasive capabilities of DRCs. CD47, often referred to as the “don’t eat me” signal (34), prevents phagocytosis by immune cells, and its elevated levels suggest a strategic adaptation by DRCs to evade immune surveillance, thereby enhancing their survival in the circulatory system and metastatic niches. FAT1 is involved in modulating cell adhesion and signaling networks (35, 36); its up-regulation potentially facilitates more robust cell-cell interactions and could promote the formation of invasive structures.

**Figure 6.**
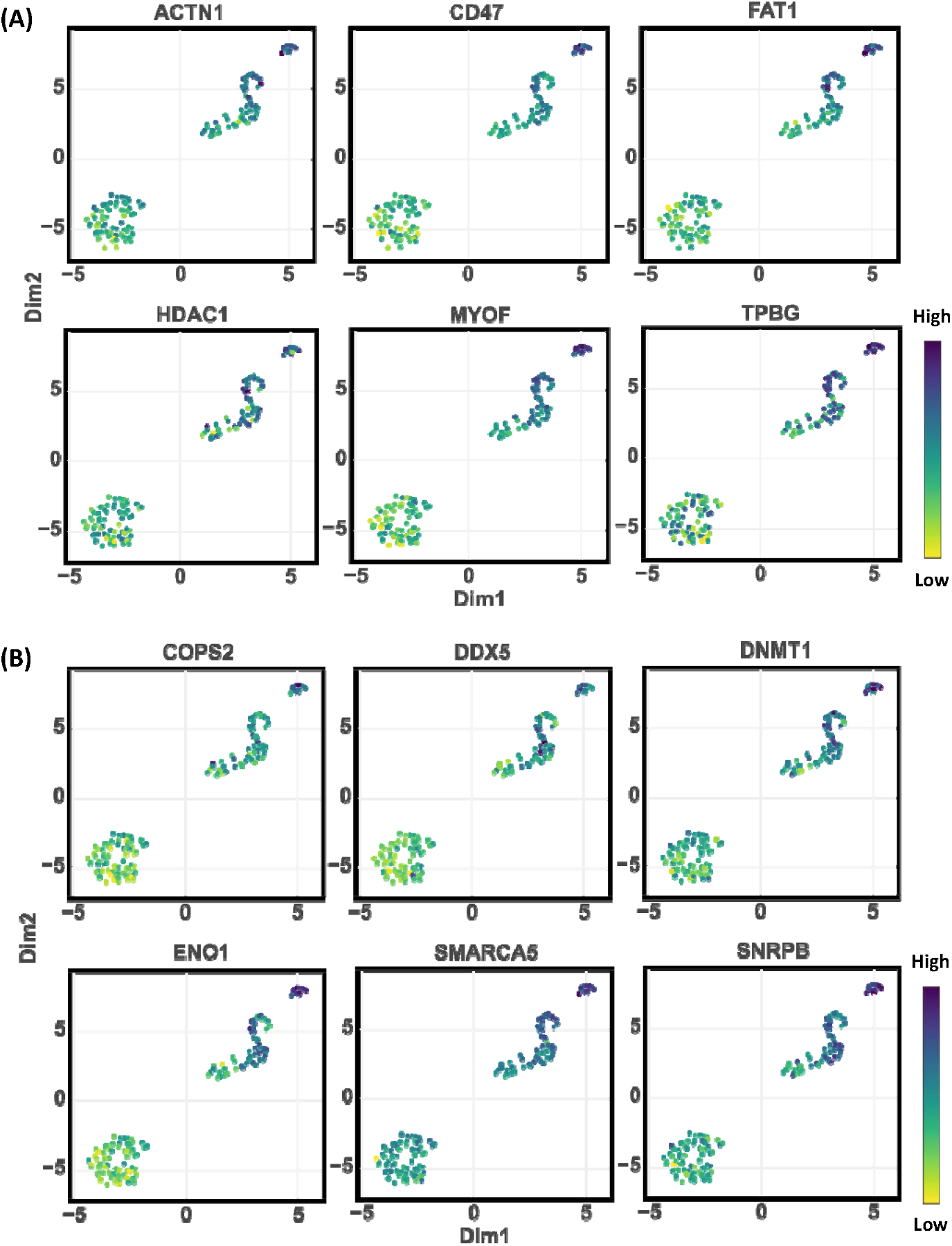
Significantly up-regulated proteins in DRCs (t-test p-value < 0.05). (A) Key surface plasma membrane proteins were selected by using the cell surface protein atlas (http://wlab.ethz.ch/cspa/). (B) Transcription factors were taken by the list (32).

Furthermore, the elevated levels of transcription factors identified, including COPS2, DDX5 (DEAD-box helicase 5), DNMT1 (DNA methyltransferase 1), ENO1, SMARCA5, and SNRPB, highlight a dynamic alteration in gene expression regulation and epigenetic modifications within DRCs (Figure 6B). These factors are integral to various cellular processes such as RNA processing, chromatin remodeling, and gene silencing. For instance, DDX5 is crucial for the processing of RNA, influencing various aspects of mRNA splicing and ribosome biogenesis (37), which are essential for rapid protein synthesis in fast-growing cancer cells. DNMT1’s role in maintaining DNA methylation patterns is critical for the epigenetic regulation of gene expression (38), suggesting that its up-regulation could support aberrant cancer cell proliferation by silencing tumor suppressor genes and activating oncogenic pathways.

The collective up-regulation of these proteins and factors in DRCs suggests a complex reprogramming of cellular functions that not only enhances the aggressive characteristics of these cells, such as invasiveness and immune evasion, but also supports a robust infrastructure for sustained growth and survival under adverse conditions. This adaptation likely contributes to the formidable challenge DRCs present in clinical settings, underscoring the need for targeted therapies that can disrupt these specific molecular mechanisms.

### Comparison of SCP and scRNA-seq in Drug-Resistant Cells

To further understand the molecular changes in drug-treated cells, we compared the results of single-cell proteomics (SCP) with RNA-seq data (Supplementary table 4 and 5). This integrative analysis revealed consistent changes across both omics platforms, providing a comprehensive view of gene expression and protein abundance alterations in drug-treated cells. We identified six genes that were upregulated and ten genes that were downregulated in both the SCP and RNA-seq datasets (Figure 7), indicating a robust regulatory pattern in drug-treated cells. LIMA1, DHRS1, S100A13, EDF1, UBE2H, and EPN1 showed increased expression levels at both the transcriptomic and proteomic levels, suggesting these molecules play a crucial role in the cellular adaptation to drug resistance. Notably, LIMA1 (LIM domain and actin-binding protein 1) is involved in actin cytoskeleton organization, which may contribute to the enhanced motility and invasiveness of drug-resistant cells. Similarly, S100A13 is associated with stress responses and extracellular signaling, further underscoring the aggressive nature of these cells. MCM7, MTMR2, HSPE1, MDH1, ASPH, HDGF, ACSL3, CSDE1, ERI3, and AHCYL1 were significantly downregulated in both omics datasets. MCM7, a member of the minichromosome maintenance complex, plays a critical role in DNA replication initiation, and its downregulation may suggest a slower replication cycle in drug-treated cells. The downregulation of proteins involved in metabolic processes, such as MDH1 (malate dehydrogenase 1) and ASPH (aspartate-β-hydroxylase), could indicate alterations in the energy production pathways, which may contribute to the survival of these cells under drug stress.

**Figure 7.**
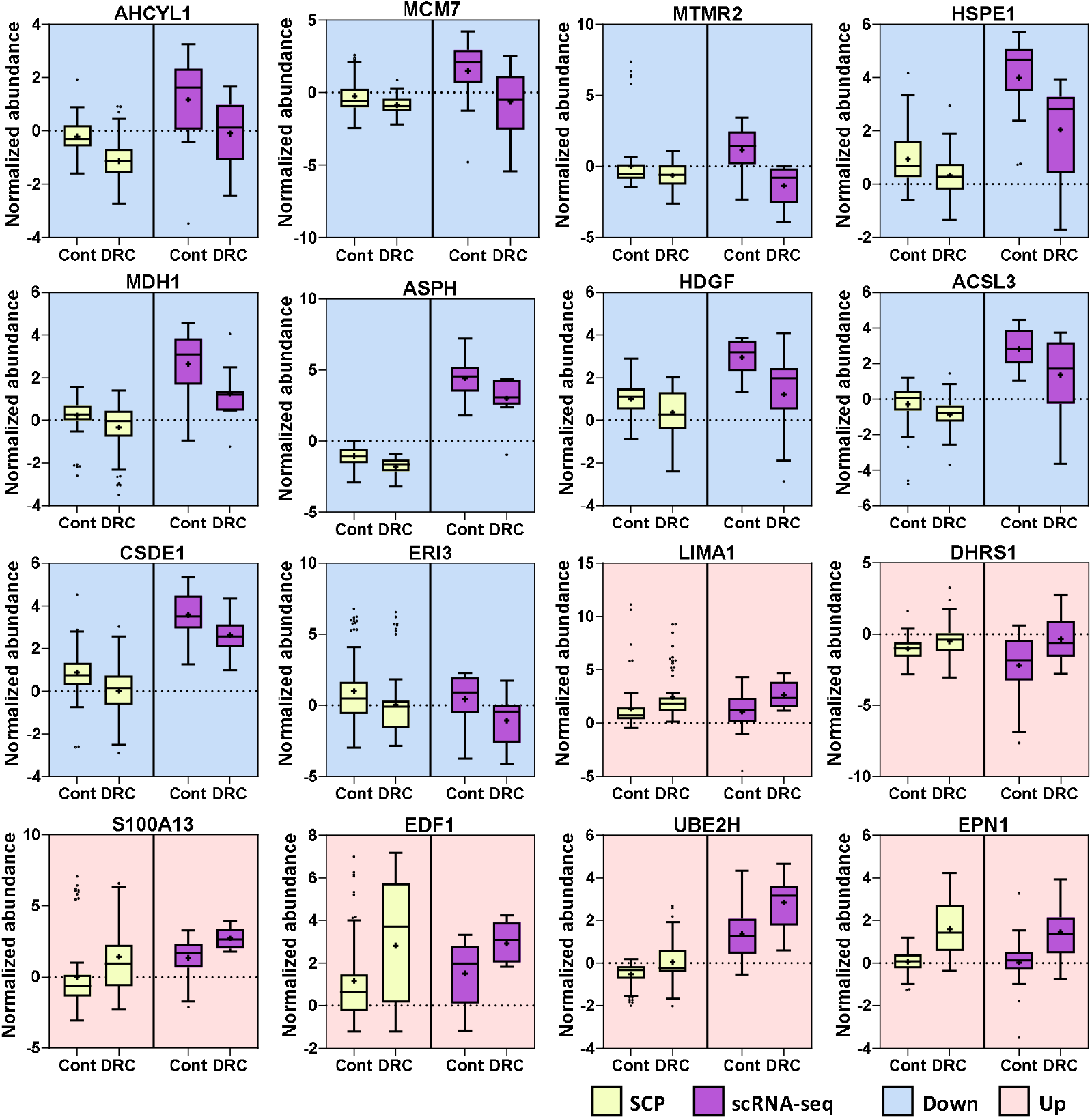
Genes and proteins significantly altered in both single-cell proteomics (SCP) and scRNA-seq datasets. The box plots represent significantly regulated genes/proteins commonly identified in both SCP and scRNA-seq analyses, with a t-test p-value < 0.05. The purple boxes correspond to the gene expression data, while the yellow boxes represent protein expression data. Each gene/protein pair is displayed side by side, illustrating the congruence between transcriptomic and proteomic regulation across drug-resistant and parental cell lines.

## Discussion

An important aspect of this study was the evaluation of two different mass spectrometry (MS) instruments for single-cell proteomics: the TimsTOF HT and the Orbitrap Ascend. Both instruments were tested for their capability to identify and quantify proteins at the single-cell level, and their performance was compared based on factors such as sensitivity, reproducibility, and depth of coverage. While both instruments demonstrated strong performance, we ultimately chose the Orbitrap Ascend coupled with Field Asymmetric Ion Mobility Spectrometry (FAIMS) for our single-cell proteomic analysis. The decision was based on the superior ability of this instrument to handle low-input samples, its enhanced signal-to-noise ratio, and its consistent detection of low-abundance proteins. The FAIMS feature of the Orbitrap Ascend significantly reduced chemical noise and improved the detection of peptides, making it particularly advantageous for single-cell proteomics where sample input is limited. Additionally, the Orbitrap Ascend’s ability to dynamically adjust ion accumulation through Automatic Gain Control (AGC) allowed for stable and reproducible measurements, further contributing to the high quality of the proteomic data generated. This consistency was critical in identifying distinct proteomic sub-clusters within the drug-resistant cell (DRC) population, which may have been more challenging to detect with other MS platforms.

The single-cell proteomic analysis of drug induced DRCs and the parental prostate cancer cells revealed a substantial increase in both the number of identified proteins and total protein abundance in DRCs compared to parental PC3 cells. The strong correlation between cell volume and proteome abundance underscores the importance of cell size in understanding the proteomic capacity and functionality of DRCs. Larger DRCs, with their increased protein content, may have a higher metabolic and biosynthetic activity, contributing to their survival and adaptability under therapeutic stress.

Unsupervised clustering analysis demonstrated clear proteomic distinctions between parental PC3 cells and DRCs, with DRCs further subdivided into three sub-clusters. This intra-population heterogeneity, associated with cell size, suggests that DRCs may comprise multiple subtypes with potentially different biological characterization. Such size-related diversity could have profound implications for understanding the biology of cancer cell survival, proliferation, and metastasis, especially in the context of therapeutic resistance. These insights not only emphasize the necessity for single-cell resolution in studying cancer heterogeneity but also highlight how advanced proteomic profiling can illuminate the complex interplay between cellular morphology and molecular function. This enhanced understanding could pave the way for more targeted and effective therapeutic strategies that consider both the molecular and morphological diversity within tumorous cell populations. Understanding these sub-clusters’ specific characteristics could inform the development of targeted therapies aimed at exploiting the unique weaknesses of each DRC subtype.

The identification of unique molecular signatures in DRCs provides insights into the pathways and processes that differentiate them from parental PC3 cells. Down-regulated proteins such as AHCYL1, APLP2, and CTNN1D in DRCs may indicate the suppression of pathways involved in normal cellular functions, impacting cell adhesion and protein synthesis. Conversely, up-regulated surface plasma membrane proteins like ACTN1, CD47, and FAT1 suggest enhanced pathways related to cell structure, immune evasion, and cell adhesion. Additionally, transcription factors such as COPS2, DDX5, and DNMT1 were up-regulated, indicating potential changes in gene expression regulation that may drive the unique characteristics of DRCs. The integration of SCP and RNA-seq data provided a robust validation of these changes at both the proteomic and transcriptomic levels. These proteomic changes, in combination with the transcriptomic data, suggest that drug-treated prostate cancer cells employ a dual strategy for survival: increasing their cellular flexibility and invasive potential while downregulating key metabolic and proliferative pathways. This reprogramming enables them to withstand therapeutic pressure and may explain their enhanced ability to evade chemotherapy and establish metastases. Targeting these altered pathways, particularly those involved in cytoskeletal dynamics, immune evasion (e.g., CD47), and metabolism, may provide new opportunities for therapeutic intervention.

This study’s findings highlight the potential of single-cell proteomics in providing detailed molecular insights into the heterogeneity and complexity of cancer cells. By combining these detailed proteomic insights with advanced single-cell analysis techniques, we aim to better understand the molecular underpinnings of drug-resistant cancer cell behavior and develop targeted therapeutic strategies to combat their aggressive and resistant nature. This study serves as a foundational step towards more personalized and effective cancer treatments, highlighting the critical role of single-cell proteomics in cancer research.

## Supporting Information

Table S1: Identified proteins and their log2-transformed abundances, Table S2: Significantly expressed proteins among the clusters, Table S3: Significantly regulated proteins between parental PC3 cells and drug-resistant cells, Table S4: Quantified proteins (normalized and imputed), Table S5: Quantified transcript genes (normalized and imputed).

## Data availability

The mass spectrometry proteomics data have been deposited to the ProteomeXchange Consortium (39) via the PRIDE partner repository with the dataset identifier PXD056872.

## ACKNOWLEDGMENTS

This work was supported by National Institutes of Health, National Cancer Institute, the Clinical Proteomic Tumor Analysis Consortium (CPTAC, U24CA271079), the Early Detection Research Network (EDRN, U2CCA271895), and Pancreatic Cancer Detection Consortium (PCDC, U01CA274514).

